# Thioredoxin reductase controls the capacity of peroxiredoxins to limit mitochondrial H_2_O_2_ release

**DOI:** 10.1101/2021.07.08.451718

**Authors:** Michaela Nicole Hoehne, Lianne J.H.C. Jacobs, Kim Jasmin Lapacz, Lena Maria Murschall, Teresa Marker, Bruce Morgan, Mark Fricker, Vsevolod V. Belousov, Jan Riemer

**Affiliations:** Department for Chemistry, Institute for Biochemistry, Redox Biochemistry, University of Cologne, Zuelpicher Str. 47a, 50674 Cologne, Germany; Institute of Biochemistry, Centre for Human and Molecular Biology (ZHMB), Saarland University, Saarbruecken, Germany; Department of Plant Sciences, University of Oxford, South Parks Road, Oxford, OX1 3RB, UK; Department of Metabolism and Redox Biology, Shemyakin-Ovchinnikov Institute of Bioorganic Chemistry, Moscow 117997, Russia; Center for Precision Genome Editing and Genetic Technologies for Biomedicine, Pirogov Russian National Research Medical University, Moscow 117997, Russia; Federal Center of Brain Research and Neurotechnologies, FMBA, Moscow 117997, Russia; Institute for Cardiovascular Physiology, Georg August University Göttingen, Göttingen 37073, Germany; Cologne Excellence Cluster on Cellular Stress Responses in Aging-Associated Diseases (CECAD), University of Cologne, 50931 Cologne, Germany

**Author notes:** correspondence to J.R., lead contact.

**Keywords:** HyPer7, mitochondria, hydrogen peroxide release, peroxiredoxin

## Abstract

H_2_O_2_ performs central roles in signaling at physiological levels, while at elevated levels it causes molecular damage. Mitochondria are major producers of H_2_O_2_, which has been implied in regulating diverse processes inside and outside the organelle. However, it still remains unclear whether and how mitochondria in intact cells release H_2_O_2_. Here we employed the genetically encoded high-affinity H_2_O_2_ sensor HyPer7 in mammalian tissue culture cells to investigate different modes of mitochondrial H_2_O_2_ release. We found substantial heterogeneity of HyPer7 dynamics between individual cells, and observed H_2_O_2_ released from mitochondria directly at the surface of the organelle and in the bulk cytosol, but not in the nucleus nor on the plasma membrane, pointing to steep gradients emanating from mitochondria. These gradients are controlled by cytosolic peroxiredoxins that act redundantly and are present with a substantial reserve capacity. Furthermore, dynamic adaptation of cytosolic thioredoxin reductase levels during metabolic changes results in improved H_2_O_2_ handling and explains previously observed cell-to-cell differences. Thus, our data indicate that H_2_O_2_-mediated signaling likely occurs close to mitochondria during specific metabolic conditions.

**HIGHLIGHTS:**

- Mitochondrial H_2_O_2_ can be detected in the cytosol in intact human cells
- Mitochondrial H_2_O_2_ gradients are steep and controlled by peroxiredoxins 1 and 2
- Peroxiredoxins 1 and 2 complement for each other
- Peroxiredoxins 1 and 2 are present with a substantial reserve capacity
- Metabolism-induced changes of reducing processes control peroxiredoxin activity

## INTRODUCTION

Reactive oxygen species (ROS) are toxic but also function as signaling molecules (Brand, 2020, D’Autreaux & Toledano, 2007, Holmstrom & Finkel, 2014, Janssen-Heininger, Mossman et al., 2008, Milev, Rhee et al., 2018, Rhee, 1999, Riemer, Schwarzlander et al., 2015, Schieber & Chandel, 2014, Sies & Jones, 2020, Winterbourn, 2020). Studies measuring ROS, in particular hydrogen peroxide (H_2_O_2_) release from isolated mitochondria as well as extracellular H_2_O_2_ in cell culture, have indicated that mitochondria are major contributors to cellular H_2_O_2_ production (Brand, 2010, Brand, 2016, Diebold & Chandel, 2016, Drose & Brandt, 2012, Klimova & Chandel, 2008, McManus, Murphy et al., 2014, Murphy, 2009, Wong, Benoit et al., 2019). Thus, not surprisingly, elevated mitochondrial H_2_O_2_ levels have been implied to drive a wide range of physiological responses and pathologies. These include responses in apoptosis, autophagy, cellular senescence, and HIF1α signaling, as well as roles in cell proliferation, migration, differentiation, and cell cycle progression (reviewed in e.g. (Brand, 2020, Chandel, 2014, Sies & Jones, 2020)).

The mitochondrial respiratory chain and associated substrate dehydrogenases are the major generators of mitochondrial ROS, with respiratory chain complexes I and III as the main sites (Brand, 2010, Murphy, 2009). This makes these sites central to physiological and presumably pathological effects caused by mitochondrial ROS. Prominent examples linking ROS generation at these sites with physiological outcomes include hypoxia signaling, where the induction of complex III-dependent ROS generation through antimycin A treatment impacted cytosolic HIF1α stabilization (Klimova & Chandel, 2008), and ischemic reperfusion injury, where reverse electron flow through complex I and consequent ROS generation underlies the pathologic consequences (Chouchani, Pell et al., 2016, Martin, Costa et al., 2019). Complex I and III release ROS to different sides of the mitochondrial inner membrane (IMM), complex I towards the matrix and complex III towards the intermembrane space (IMS) (Brand, 2010, Han, Williams et al., 2001) (although release of ROS from complex III to the matrix has also been reported (Muller, Liu et al., 2004)). This spatial organization of ROS generation and release has been confirmed by redox proteomics data on isolated mitochondria. When ROS production was induced by inhibitor treatment of complexes I and III, proteins that were oxidatively modified localized mainly to matrix and IMS, respectively (Bleier, Wittig et al., 2015).

The proximal ROS produced by these complexes are superoxide anions that are rapidly dismutated to H_2_O_2_ and oxygen either by superoxide dismutase 2 (SOD2) in the matrix, or SOD1 in the IMS depending on the site of their generation. H_2_O_2_ (but not superoxide) can diffuse (slowly) across the IMM, while porins/VDACs in the outer membrane (OMM) likely allow membrane transfer with little interference. Also, for the IMM, transport might be facilitated by a yet unknown transporter as is the case for other cellular membranes (Bienert, Moller et al., 2007, Calamita, Ferri et al., 2005, Marchissio, Frances et al., 2012). Compartmental concentrations of H_2_O_2_ are set mainly by rates of production (activities of superoxide generator sites and SODs), removal through the activities of antioxidative enzyme systems (peroxiredoxins, catalases), diffusion of H_2_O_2_ out of or into the compartment of interest (*e.g.* mitochondrial H_2_O_2_ release to the cytosol), or side reactions with biomolecules like proteins or lipids. Thus, to act as signaling molecule outside mitochondria, mitochondrial H_2_O_2_ has to be produced in sufficient amounts, it has to escape “premature handling” by local antioxidative systems or side reactions with biomolecules, it has to navigate the complex morphology of mitochondria to reach the cytosol, and it has to compete with potent cytosolic antioxidative systems for its target molecules, such as redox-regulated proteins.

The dynamics of H_2_O_2_ inside mitochondria and its release from mitochondria is only poorly understood in mammalian cells. Previous studies relied on different tools including matrix-targeted or untargeted chemical probes (*e.g.* Mito-SOX) or genetically encoded probes with low sensitivity (*e.g.* HyPer3 or roGFP2-Orp1). In addition, H_2_O_2_ release from isolated mitochondria or increased extracellular H_2_O_2_ has been reported (Goncalves, Quinlan et al., 2015, Kalinovic, Oelze et al., 2019, Liao, Franco-Iborra et al., 2020, Plecita-Hlavata, Engstova et al., 2020, Roma, Deponte et al., 2018). The chemical probes do not allow high resolution monitoring of extramitochondrial H_2_O_2_, and lack dynamic reversibility (*i.e.* it is difficult to assess dynamic fluxes of H_2_O_2_), while low sensitivity genetically encoded probes only respond to strong exogenous H_2_O_2_ treatments or report on the glutathione pool required to re-reduce the sensor, rather than directly on H_2_O_2_ levels. Furthermore, measurements on isolated mitochondria necessarily ignore the role of cytosolic redox systems and thus fail to represent the cellular situation. Likewise, measurements outside cells with highly sensitive methods might detect concentrations well below the point of biological activity, but fail to directly report on mitochondrial H_2_O_2_ release and the local internal concentrations or gradients. Thus, although mitochondrial H_2_O_2_ appears to have important roles in many signaling or pathological phenotypes, *in situ* evidence for its release from mitochondria in intact cells and its subsequent dynamics in the cytosol remains scarce.

Recently, a high-affinity, pH-insensitive H_2_O_2_ sensor, HyPer7, was introduced that allows relatively rapid and reversible dynamic measurements of highly localized H_2_O_2_ concentrations (Pak, Ezerina et al., 2020). Notably, experiments with HyPer7 cast doubt on the release of H_2_O_2_ from mitochondria (Pak et al., 2020). Instead, they indicated that only upon inhibition of the cytosolic thioredoxin system, could mitochondrial H_2_O_2_ diffuse into the cytosol at a measurable amount. Here, we employed the HyPer7 sensor to revisit mitochondrial H_2_O_2_ release. We demonstrate that in different settings mitochondria do indeed release sufficient H_2_O_2_ to be detectable on the OMM and in the bulk cytosol. The release is strictly controlled by the metabolic state of the cell in a janus-faced manner: on the one hand higher activity of the respiratory chain in cells grown on galactose increases H_2_O_2_ generation, on the other hand it also increases the levels of the cytosolic anti-oxidant thioredoxin system, which limits efficient functioning of the cytosolic two-Cys peroxiredoxins PRDX1 and PRDX2. Cells appear to contain a reserve capacity of these peroxiredoxins which act redundantly. Our results thus indicate synchronized adaptation of redox systems in the cytosol in response to changing metabolic states that influence mitochondrial H_2_O_2_ release, whilst still allowing direct H_2_O_2_ signaling in close proximity to mitochondria.

## RESULTS

### Antimycin A treatment induces cytosolic responses of HyPer7

Little is known about the spatio-temporal organization of H_2_O_2_ release or removal in intact human cells. Thus, we sought to characterize H_2_O_2_ fluxes originating from mitochondria as the predominant source of intracellular H_2_O_2_. To this end, we made use of the genetically encoded fluorescent probe, HyPer7, which allows the real-time monitoring of basal H_2_O_2_ levels with unprecedented sensitivity in specific subcellular compartments ((Pak et al., 2020); **Figure 1A**). HyPer7 comprises a circular permutated yellow fluorescent protein (cpYFP) genetically inserted into the OxyR regulatory domain (OxyR-RD) of *Neisseria meningitis*. The OxyR-RD moiety of the HyPer7 probe responds directly to H_2_O_2_ by forming a disulfide bond. This in turn induces a shift in the excitation spectrum of the cpYFP. While the probe is oxidized by H_2_O_2_, it is predominantly re-reduced by endogenous thioredoxins (Kritsiligkou, Shen et al., 2021). HyPer7 oxidation is therefore determined by rapid H_2_O_2_-driven oxidation and slower thioredoxin-driven reduction (**Figure 1A**, (Pak et al., 2020)). In our experimental setup, the readout of HyPer7 can be presented as ratio of the excitation wavelengths at 469 and 390 nm that automatically compensate for varying probe concentrations and to ensure pH independence of the measurements (for data acquisition and analysis using a multi-well microscope setup, see Materials and Methods). High ratios indicate oxidized and low ratios reduced OxyR-RD moiety, respectively. This was clearly visible when we monitored single cell responses of cytosolic HyPer7 to the repeated addition of exogenous H_2_O_2_ in HEK293 cells (**Figure 1B**). We observed that with amounts of external H_2_O_2_ as low as 2 μM, HyPer7 became oxidized, however to a markedly different degree in individual cells. To transparently report on this cell-to-cell heterogeneity that we observed in all our experiments with HyPer7, while still allowing for an easy assessment of the data, we present our data both as average over many cells (**Figure 1A** and **B**, solid black line) and as single cell data (**Figure 1A** and **B**, light gray data points).

**Figure 1.**
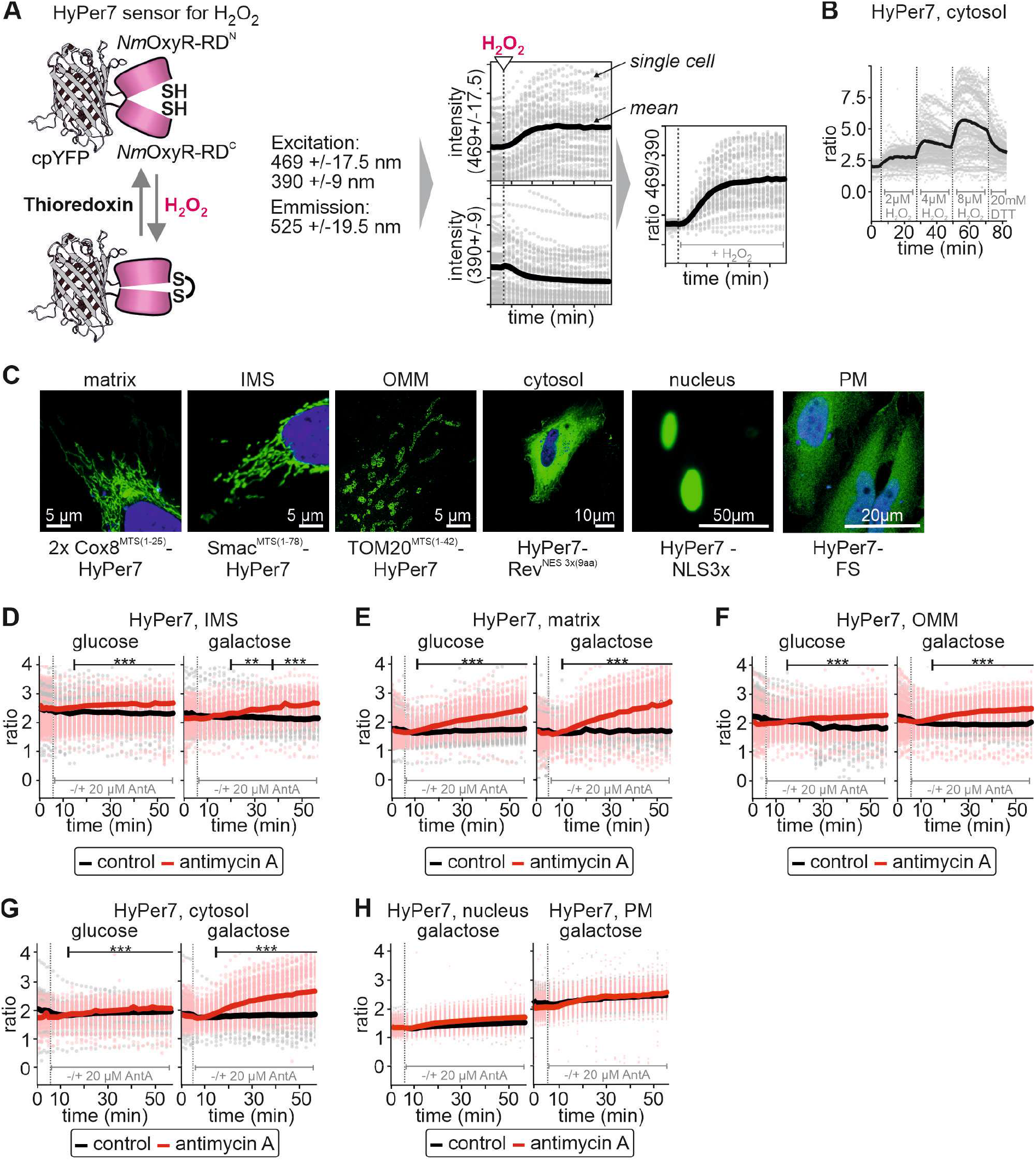
Antimycin A treatment induces responses of cytosolic HyPer7. **A.** The H_2_O_2_ sensing mechanism of the HyPer7 probe and representation of data. **B.** The response of cytosolic HyPer7 in HEK293 cells to repeated bolus of exogenous H_2_O_2_ at the indicated concentrations. Solid line represents mean, light gray points are ratios measured in individual cells. **C.** Confirmation of sensor localization to different cellular compartments. Green, HyPer7; blue, DAPI stain **D.-H.** Response of HyPer7 probes targeted to indicated compartments to incubation with antimycin A (a complex III inhibitor, red curve and data points) or ethanol as control (black line and data points). HEK293 cells were grown either with glucose or galactose as carbon source as indicated. Solid line represents mean, points colored in the lighter version of the respective color are the corresponding ratios measured in individual cells. The numbers of cells per experiment as well as information on the statistical analysis for each dataset can be found in supplemental table **S6**. **, p ≤ 0.01, ***, p ≤ 0.001

To obtain a dynamic picture of the cellular H_2_O_2_ landscape upon induction of H_2_O_2_ production in mitochondria, we also targeted HyPer7 to the mitochondrial matrix, IMS, the cytosolic side of the OMM, the plasma membrane (PM), and the nucleus (**Figure 1C**). We then monitored the probe response upon incubation of HEK293 cells with the complex III inhibitor antimycin A. Antimycin A causes release of superoxide anions (which are rapidly dismutated to H_2_O_2_) towards the IMS side of the IMM. We performed these experiments with cells grown in glucose or galactose (cells were adapted to galactose growth for at least one week) as the carbon source. Compared to glucose, galactose enhances oxidative metabolism and thus in principle upon antimycin A treatment should generate more superoxide anions. In the IMS, we observed a small HyPer7 response that was more pronounced in cells grown on galactose than in cells grown on glucose (**Figure 1D**). Interestingly, matrix-targeted HyPer7 appeared to react even more strongly than its IMS counterpart, likely representing diffusion of H_2_O_2_ over the IMM from the cristae space or partial direct release of ROS from complex III towards the matrix (**Figure 1E**). Similar to the IMS and matrix, we also observed a stronger response in cells grown on galactose for the OMM and cytosolic HyPer7 (**Figures 1F,G**). Notably, cytosolic HyPer7 reacted only to antimycin A treatment when cells were grown on galactose. However, no Hyper7 response was observed to antimycin A treatment in the nucleus or on the PM, in cells grown on galactose (**Figure 1H**). In summary, antimycin A treatment resulted in detectable deflections of HyPer7 at the OMM and in the cytosol, but not in the nucleus or at the plasma membrane. HyPer7 responses in cells grown in galactose-containing medium were in general more pronounced, indicating that the steep H_2_O_2_ gradient emanating from mitochondria is modulated by metabolic adaptations (*e.g.* differences in H_2_O_2_ generation).

### Galactose-grown cells present stronger HyPer7 responses upon reoxygenation after hypoxia

Next, we assessed this metabolic effect in a different experimental regime. To this end, we exposed cells to hypoxia and subsequent reoxygenation after ca. 200 min, or to continuous hypoxia. Reoxygenation should increase the generation of H_2_O_2_ particularly in mitochondria. During hypoxia, the absolute HyPer7 ratios in all compartments assessed was lower than in normoxic cells, but remained unchanged over an extended period indicating that hypoxia alone does not result in substantial H_2_O_2_ generation (**Figures 2A-D,** black line and red line before 200 min). Upon reoxygenation HyPer7 became oxidized in all compartments assessed, particularly if cells had been grown in galactose-containing medium (**Figures 2A-D**). In cells maintained constantly in hypoxic conditions HyPer7 ratios did not change (**Figures 2A-D**, black line).

**Figure 2.**
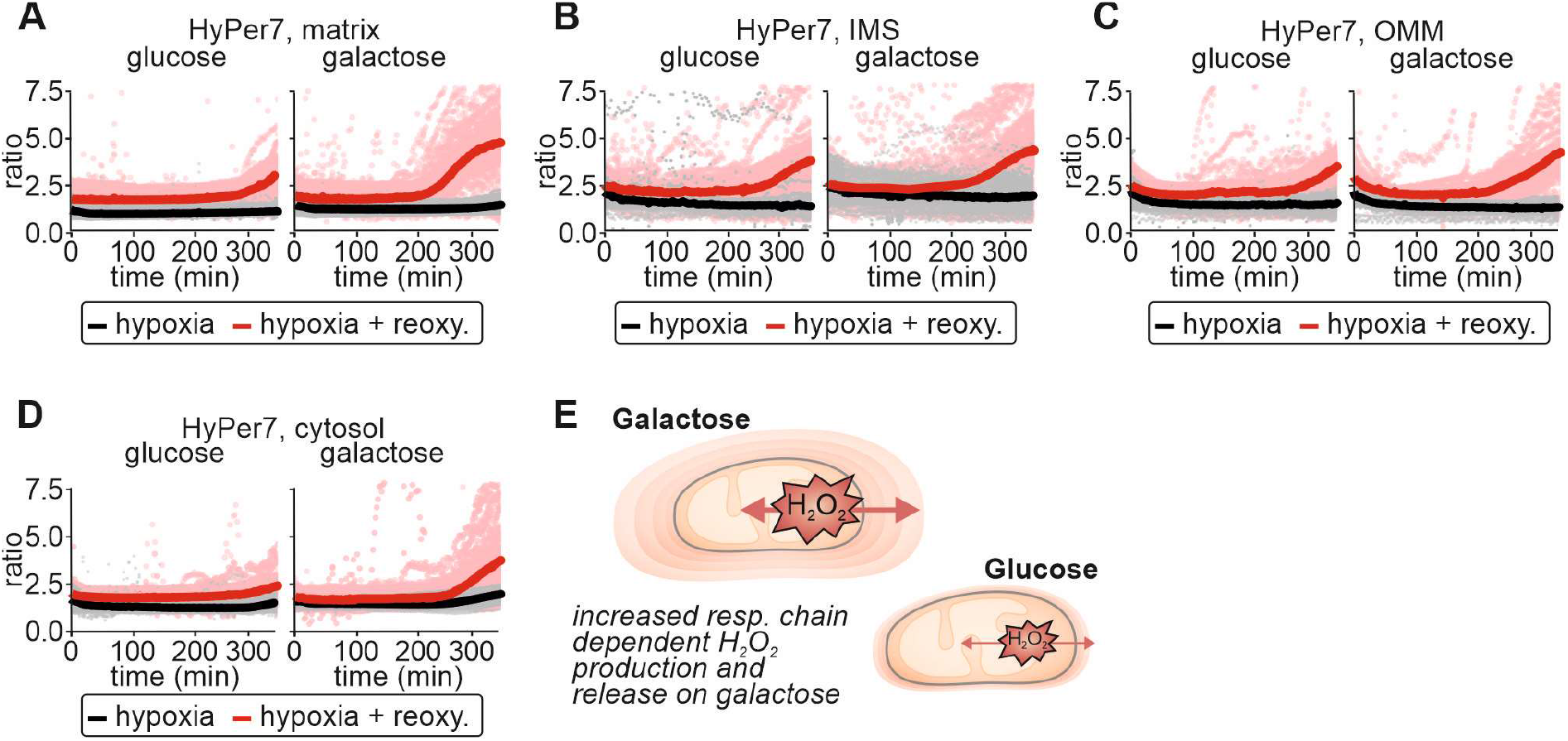
Galactose-grown cells present with stronger HyPer7 responses upon reoxygenation after hypoxia. **A-D.** Response of HyPer7 probes targeted to indicated compartments to reoxygenation after hypoxia (red curve and data points) or to continued hypoxia (black line and data points). HEK293 cells were grown either with glucose or galactose as carbon source. Solid line represents mean, points colored in the lighter version of the respective color are the corresponding ratios measured in individual cells. **E.** Model. Cells grown on galactose-containing medium exhibit increased production of H_2_O_2_ and thus also increased detection of cytosolic H_2_O_2_ indicating H_2_O_2_ gradients of different steepness around mitochondria. The numbers of cells per experiment as well as information on the statistical analysis for each dataset can be found in supplemental table **S6**.

Hypoxia caused a reduction in the steady-state ratios in all compartments assessed. However, reoxygenation after hypoxia lead to substantial oxidation of HyPer7, particularly in galactose-grown cells, emphasizing the importance of metabolic conditions for the generation and handling of H_2_O_2_ (**Figure 2E**).

### Equal mitochondrial H_2_O_2_ generation results in detection of different cytosolic H_2_O_2_ amounts upon growth in the presence of different carbon sources

The amounts of H_2_O_2_ generated through incubation of cells with antimycin A depend on the activity of the respiratory chain. We modulated electron flux through the respiratory chain by providing cells with different carbons sources. However, this also impacts many other cellular (including antioxidative) processes, *e.g.* flux through the pentose phosphate pathway. To distinguish the effects of increased mitochondrial H_2_O_2_ production from general changes in cellular processes in cells grown on galactose, we turned to a genetically engineered H_2_O_2_ producing system using matrix-targeted C-terminally FLAG-tagged D-amino acid oxidase (mtDAO, **Figure 3A**, (Matlashov, Belousov et al., 2014, Pak et al., 2020)). Using the TRex-FlpIn system, we generated stable cell lines to ensure homogenous expression of mtDAO in all cells. Moreover, mtDAO expression is doxycycline-inducible to avoid adaptation effects that might result from a continuous expression of this enzyme. After induction of mtDAO expression for 24 hours, and upon addition of D-alanine but not L-alanine, mtDAO produced H_2_O_2_ in a dose-dependent manner in the mitochondrial matrix (**Figure 3B**).

**Figure 3.**
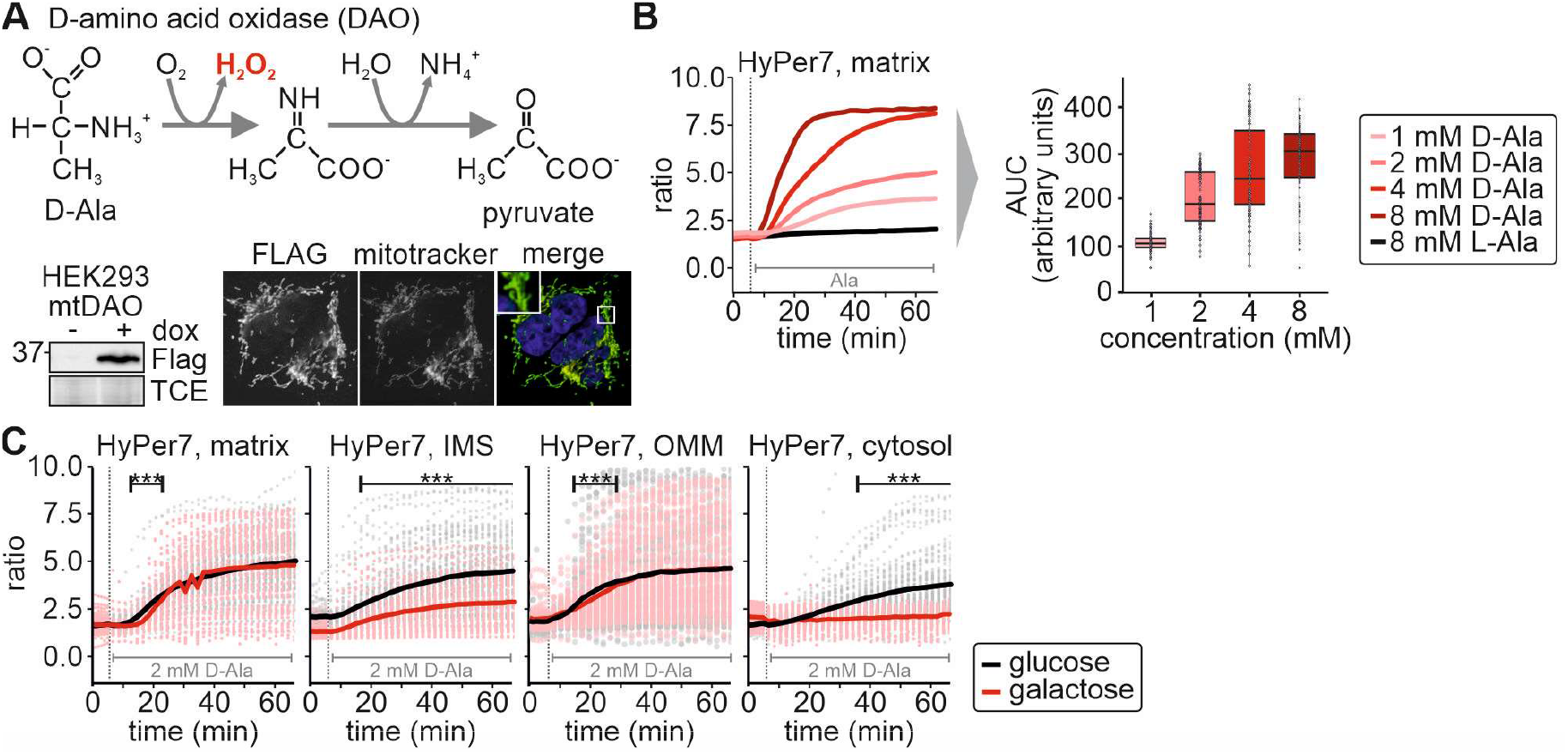
Equal mitochondrial H_2_O_2_ generation results in detection of different cytosolic H_2_O_2_ amounts upon growth in the presence of different carbon sources. **A.** Mechanism of D-amino acid oxidase (DAO) in the production of H_2_O_2_. Cell lines stably and inducibly expressing a mitochondrial matrix version of DAO (mtDAO) were generated and tested by immunoblot and immunofluorescence. **B.** Titration of D-alanine (D-Ala) in mtDAO-containing cells and monitoring by matrix HyPer7 (cell generated with the Flp-In T-REX-system). Solid line represents average of multiple measure cells. For area under the curve (AUC) analyses, means of control curves were subtracted from means of D-Ala-treated cells. **C.** Response of HyPer7 probes targeted to indicated compartments to treatment of mtDAO-expressing HEK293 cells with 2 mM D-Ala (cells generated with the Flp-In T-REX-system). HEK293 cells were grown either with glucose (black curve and data points) or galactose (red curve and data points) as carbon source. Solid line represents average, points colored in the lighter version of the respective color are the corresponding ratios measured in individual cells. The numbers of cells per experiment as well as information on the statistical analysis for each dataset can be found in supplemental table **S6**. ***, p ≤ 0.001

We then used this system to monitor HyPer7 responses in different compartments of cells grown on galactose- or glucose-containing medium (**Figure 3C**). We detected HyPer7 oxidation in all compartments upon addition of 2 mM D-alanine demonstrating that in this controlled setting H_2_O_2_ is produced and released from mitochondria. The HyPer7 response in the matrix was similar between glucose and galactose-grown cells. Interestingly, the response of HyPer7 in the cytosol and the IMS was less pronounced in cells grown on galactose compared to growth on glucose, the opposite trend to that observed with antimycin A or re-oxygenation.

Likewise, the response at the OMM was similarly independent of the growth medium, an observation that we currently cannot explain, as it also appears to differ from antimycin A and reoxygenation treatments. In the nucleus and on the PM, the HyPer7 probe only responded at 8 mM D-alanine (**Figure S1**). Collectively, our data indicate that the impact of releasing defined amounts of H_2_O_2_ to the cytosol is reduced in cells grown on galactose compared to cells grown on glucose.

### Growth in the presence of different carbon sources impacts compartmental H_2_O_2_ handling

With our different treatment regimes, we obtained seemingly contradicting results: antimycin A treatment and reoxygenation after hypoxia induced a more prominent oxidation of cytosolic HyPer7 on galactose-grown compared to glucose-grown cells, while for mtDAO-induced H_2_O_2_ generation this was *vice versa*. An explanation might be prior adaptation processes on galactose that strengthen cellular antioxidative systems. These are then primed to counteract the increased amounts of H_2_O_2_ produced as a consequence of increased flux through the respiratory chain. Antimycin A treatment would still lead to detection of high H_2_O_2_ production that exceed the capacity of the antioxidative systems; hence the stronger oxidation of HyPer7 with antimycin A on galactose. Conversely, the mtDAO system produces similar amounts of H_2_O_2_ on both glucose and galactose, but the more efficient antioxidative systems on galactose would attenuate the impact of its release. To further test this hypothesis, we exposed HEK293 cells to bolus treatments with exogenous H_2_O_2_. Indeed, in all compartments the HyPer7 response was attenuated in cells grown on galactose compared to glucose (**Figure 4A**).

**Figure 4.**
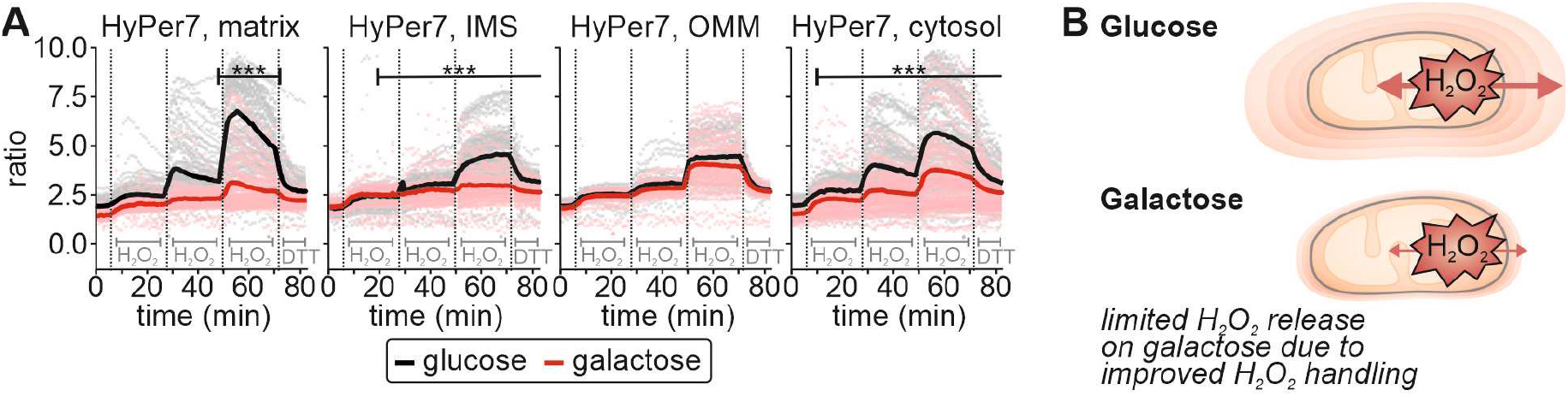
Growth in the presence of different carbon sources impacts compartmental H_2_O_2_ handling. **A.** Response of HyPer7 probes targeted to indicated compartments to treatment with increasing amounts of exogenous H_2_O_2_. HEK293 cells were grown either with glucose (black curve and data points) or galactose (red curve and data points) as carbon source. The numbers of cells per experiment as well as information on the statistical analysis for each dataset can be found in supplemental table **S6**. ***, p ≤ 0.001 **B.** Model. (Mitochondrial) H_2_O_2_ is more efficiently handled in galactose-grown cells in mitochondria and cytosol.

Thus, we demonstrate that in HEK293 cells different modes of mitochondrial H_2_O_2_ production (e.g. antimycin A, mtDAO) result in release of H_2_O_2_ from mitochondria. However, growth of cells on a carbon source that induces increased electron flux through the respiratory chain (galactose) appears to upregulate anti-oxidant pathway(s) and prepare cells to handle H_2_O_2_ more efficiently, and thus attenuates the consequences of mitochondrial H_2_O_2_ release (**Figure 4B**).

### Increased activity of the cytosolic thioredoxin system reduces the impact of mitochondrial H_2_O_2_ release during increased activity of the respiratory chain

What are the specific changes induced by continuously growing cells on galactose instead of glucose? We addressed this by performing quantitative proteomics on HEK293 cells grown on glucose or galactose (**Figure 5A, Raw data file 1**). While the amounts of many proteins differed between both conditions, only one protein was significantly altered that qualified as an antioxidative enzyme. This was cytosolic thioredoxin reductase 1, TXNRD1, which was increased about 2- to 2.5-fold in cells grown on galactose compared to glucose-grown cells. Notably, levels of peroxiredoxins remained unchanged. We confirmed this result by immunoblotting against selected redox proteins (**Figure 5B**).

**Figure 5.**
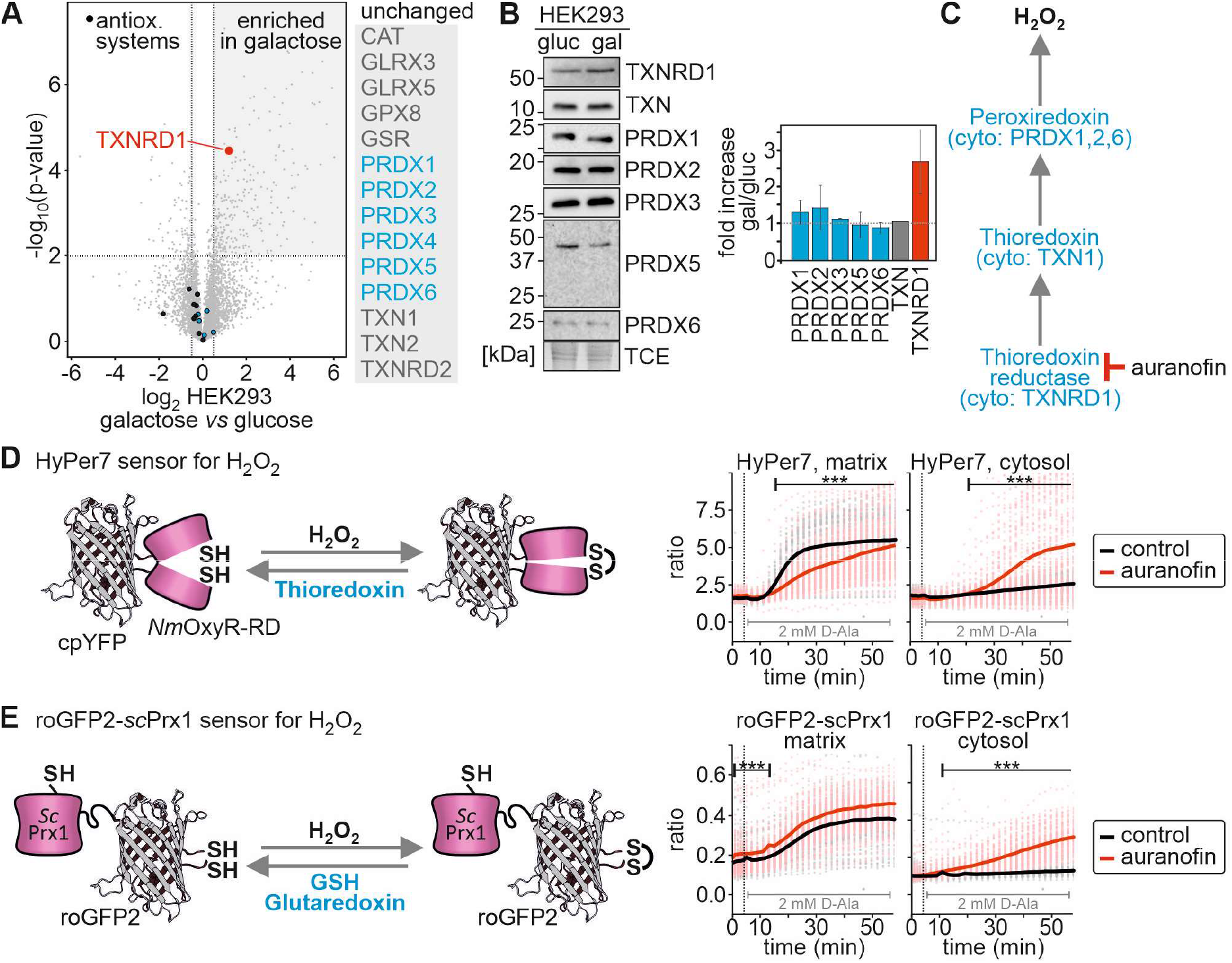
Increased activity of the cytosolic thioredoxin system limits mitochondrial H_2_O_2_ release during increased use of the respiratory chain. **A.** Protein levels in cells grown on glucose and galactose. Cell lysates were analyzed by quantitative mass spectrometry (n = 4 biological replicates). Levels of antioxidative enzymes (especially peroxiredoxins, PRDX in blue) are not changed except for cytosolic thioredoxin reductase (TXNRD1). **B.** Protein levels in cells grown on glucose and galactose. Cell lysates were analyzed by immunoblot (n = 2 biological replicates). Cytosolic thioredoxin reductase (TXNRD1) is increased by more than two-fold in cells grown on galactose compared to glucose-grown cells. **C.** Mechanisms of H_2_O_2_ degradation by peroxiredoxins. Auranofin inhibits thioredoxin reductase and thus the reductive half reaction in the detoxification of H_2_O_2_. Arrows indicate the “flux” of electrons. **D,E.** Response of HyPer7 (**D**) and roGFP2-scPrx1 (**E**) probes targeted to indicated compartments to treatment of mtDAO-expressing HEK293 cells with 2 mM D-Ala (cells generated with the Flp-In T-REX-system). HEK293 cells were incubated either with 1 μM auranofin (red curve and data points) or DMSO (black curve and data points). Solid line represents average, points colored in the lighter version of the respective color are the corresponding ratios measured in individual cells. The numbers of cells per experiment as well as information on the statistical analysis for each dataset can be found in supplemental table **S6**. ***, p ≤ 0.001

We then tested the involvement of the thioredoxin system in regulating H_2_O_2_ dynamics. To this end, we inhibited thioredoxin reductase using auranofin (**Figure 5C**). Auranofin treatment strongly increased oxidation of the cytosolic HyPer7 sensor during mtDAO-induced H_2_O_2_ generation (**Figure 5D**). The thioredoxin system affected measurements in two ways: first, by attenuating H_2_O_2_ levels in the cytosol *e.g.* by acting through peroxiredoxin, and second, by accelerating the reducing half reaction of the HyPer7 sensor (**Figure 1A,5D;**(Pak et al., 2020)). To disentangle these contributions, we employed a different H_2_O_2_ sensor that we recently developed, that is not reduced by the thioredoxin system but by the glutathione system instead (Calabrese, Peker et al., 2019, Kritsiligkou et al., 2021). It comprises a redox-sensitive green fluorescent protein (roGFP2; (Dooley, Dore et al., 2004)) genetically fused with the monothiol Prx1 from *Saccharomyces cerevisiae* (*Sc*Prx1). The *Sc*Prx1 moiety serves to efficiently transfer oxidative equivalents from H_2_O_2_ to roGFP2. This probe is predominantly reduced by endogenous GSH/glutaredoxins, which directly reduce the roGFP2 moiety and also scPrx1 itself. RoGFP2-*Sc*Prx1 oxidation is therefore determined by rapid H_2_O_2_-driven oxidation and slower GSH/glutaredoxin-driven reduction. Oxidation of this probe was also more pronounced upon auranofin treatment (**Figure 5E**) indicating that thioredoxin reductase 1 exerted its effect via lowering cytosolic H_2_O_2_ levels and to a lesser extent via accelerated probe reduction. Thus, in addition to potential metabolic adaptations (i.e. increasing the flux through the pentose phosphate pathway), it is the increased activity of the cytosolic thioredoxin system that suppresses the impact of mitochondrial H_2_O_2_ release during increased activity of the respiratory chain.

### Increased levels of cytosolic thioredoxin reductase in HeLa cells reduce the impact of mitochondrial H_2_O_2_ release to the cytosol

Our data are in contradiction to a recent study with HyPer7 that did not detect the release of H_2_O_2_ from mitochondria upon rotenone treatment and mtDAO-induced H_2_O_2_ generation, which was apparently caused by a very strict control by the cytosolic thioredoxin system (Mishina, Bogdanova et al., 2019, Pak et al., 2020). Based on the evidence so far, we predict that this is a consequence of constitutively higher levels of thioredoxin reductase in Hela cells compared to HEK293 cells. We confirmed that HeLa cells harbor higher thioredoxin reductase activity compared to HEK293 cells (**Figure 6A**, (Geiger, Wehner et al., 2012)), and also confirmed that there was no detectable H_2_O_2_ release in Hela cells, despite the fact that matrix HyPer7 becomes oxidized (**Figure 6B**). Likewise, H_2_O_2_ generation by mtDAO provoked cytosolic oxidation of HyPer7 only in HEK293, but not HeLa cells (**Figure 6C**).

**Figure 6.**
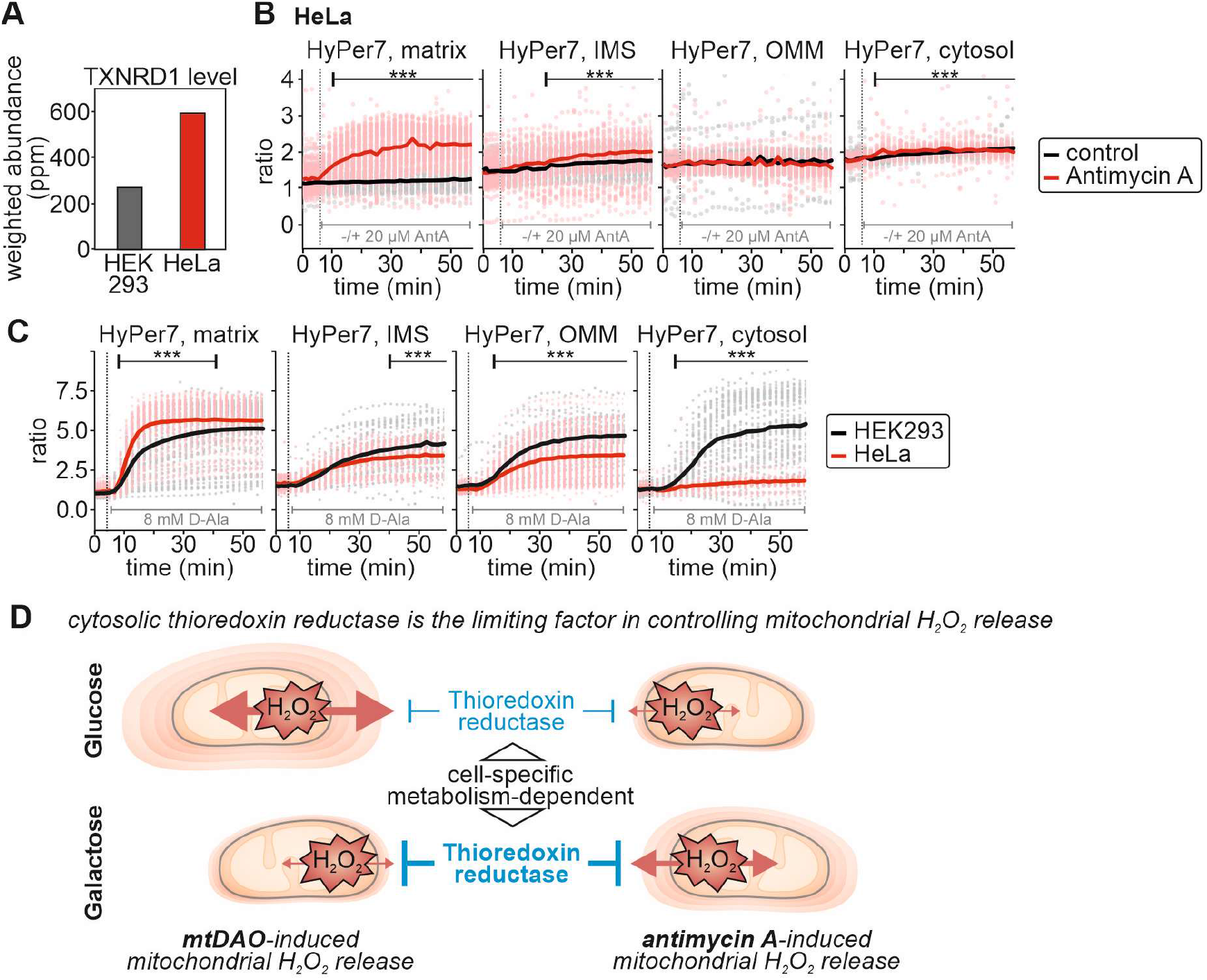
Increased levels of cytosolic thioredoxin reductase in HeLa cells prevent mitochondrial H_2_O_2_ release to the cytosol. **A.** TXNRD1 levels in HeLa and HEK293 cells. Data are from (Geiger et al., 2012). **B.** Response of HyPer7 probes targeted to indicated compartments to incubation with antimycin A (AntA, red curve and data points) or ethanol as control (black curve and data points). HeLa cells were grown with galactose as carbon source. Solid line represents average, points colored in the lighter version of the respective color are the corresponding ratios measured in individual cells. **C.** Response of HyPer7 probes targeted to indicated compartments to treatment of mtDAO-expressing HEK293 (black curve and data points; cell generated with the piggyBAC system) or HeLa (red curve and data points; cell generated with the piggyBAC system) cells with 8 mM D-Ala. Cells were grown with glucose as carbon source. Solid line represents average, points colored in the lighter version of the respective color are the corresponding ratios measured in individual cells. **D.** Model. Cells- and metabolism-specific empowerment of the thioredoxin reductase system controlling H_2_O_2_ concentrations following mitochondrial release. The numbers of cells per experiment as well as information on the statistical analysis for each dataset can be found in supplemental table **S6**. ***, p ≤ 0.001

Collectively, our data indicate that mitochondria do release H_2_O_2_, which is counteracted by cytosolic antioxidative systems driven by thioredoxin reductase. The balance between mitochondrial H_2_O_2_ release and H_2_O_2_ degradation is dynamically adapted, for example depending on metabolic conditions, but also in a cell type-specific manner. Adaptation thereby not only involves proteomic changes but likely also changes in the flux through different metabolic pathways because adaptation can take place on a time scale of minutes that is not sufficient to significantly increase thioredoxin reductase protein amounts (**Figure 6D**).

### Cytosolic peroxiredoxins cooperate in controlling mitochondrial hydrogen peroxide release

Upon growth on galactose, HEK293 cells adapted by increasing the levels of cytosolic thioredoxin reductase. This indicates that the reducing half-reaction during H_2_O_2_ detoxification by peroxiredoxins may limit H_2_O_2_ handling, rather than the levels of peroxiredoxins. Thus, we next assessed how cytosolic peroxiredoxin impacted mitochondrial H_2_O_2_ release. To this end, we generated single knockouts of the cytosolic dithiol peroxiredoxins, PRDX1 and PRDX2 using CRISPR-Cas technology (**Figure 7A,B**). In these cells, when grown on glucose, levels of other antioxidant proteins remained unchanged, in particular the concentrations of the respective other cytosolic peroxiredoxin (**Figure 7A,B**). Steady state HyPer7 ratios were also essentially unchanged in both single knockout cell lines (**Figure 7C**). Mitochondrial production of H_2_O_2_ by antimycin A treatment or in cells stably and inducibly expressing mtDAO (in these cells, stable and inducible expression of mtDAO used the piggyBAC system) revealed almost no difference between either peroxiredoxin knockout line compared to the wild type (**Figure 7D** and **S2**). Only during bolus application of exogenous H_2_O_2_, did HyPer7 in the PRDX1 and PRDX2 knockout cells exhibit an increased oxidation compared to wildtype cells, especially in matrix and cytosol (**Figure 7E**). Notably, stable and inducible expression of mtDAO with the piggyBAC system appeared to influence the HyPer7 steady state even in the absence of D-alanine (compare steady state = first 5 min between **7D** (mtDAO) and **7E** (no mtDAO). We did not observe this influence in HEK293 cells stably and inducibly expressing mtDAO using the Flp-In T-Rex system (compare to **Figure 3C**), and hypothesize that leaky expression in the piggyBAC system contributes to the increased HyPer7 steady state ratio.

**Figure 7.**
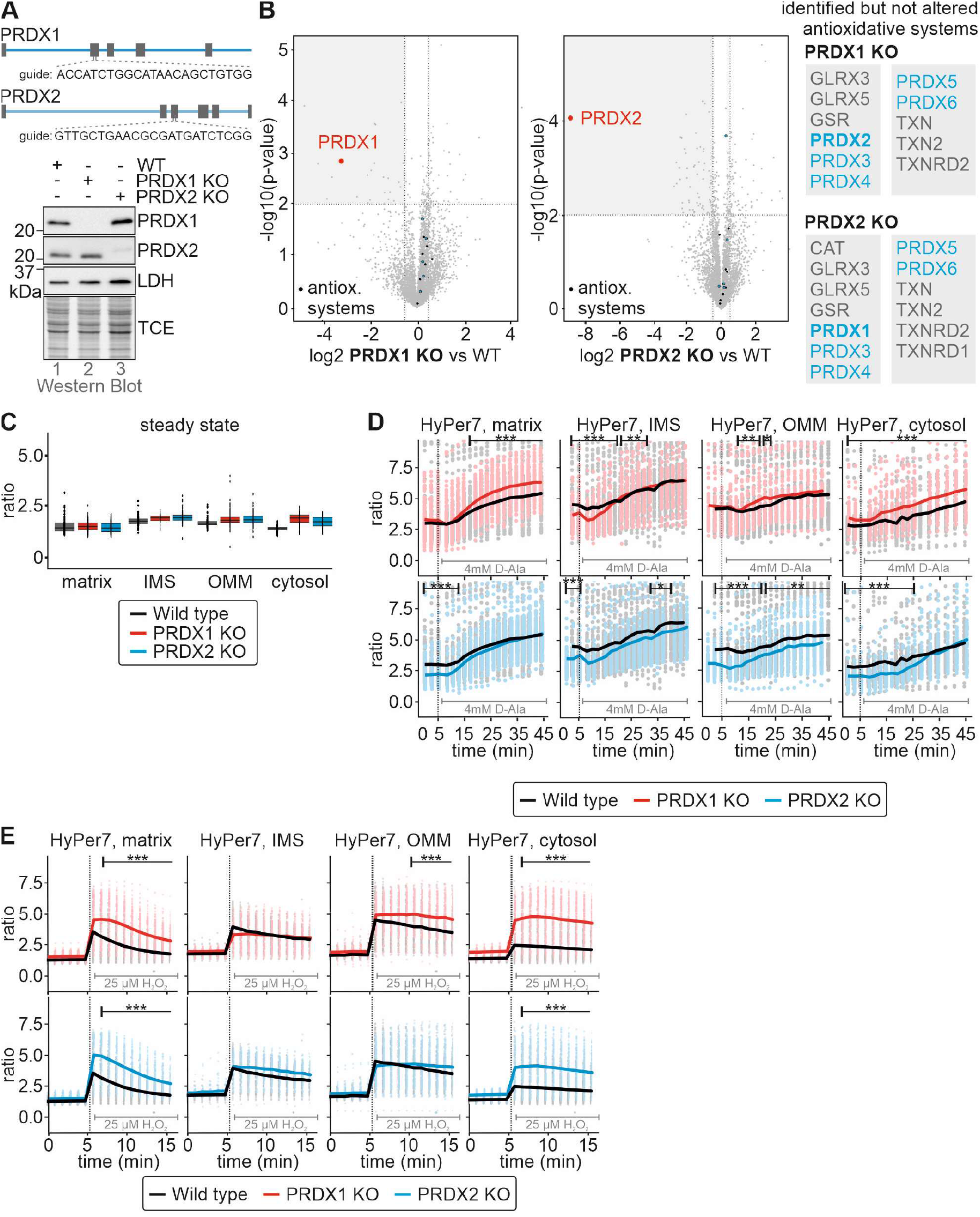
Cytosolic peroxiredoxins cooperate in controlling mitochondrial H_2_O_2_ release. **A,B.** Peroxiredoxin levels in peroxiredoxin 1 (PRDX1 KO) and peroxiredoxin 2 (PRDX2 KO) knockout cells. Lysates of the indicated cell lines grown in glucose-containing medium were analysed by immunoblot (**A**) and quantitative proteomics (n = 4) (**B**). **C.** HyPer7 steady state ratios of the indicated cell lines. HyPer7 probes were targeted to the indicated compartments of the respective cell lines. Cells were grown in glucose-containing medium. **D.** Response of HyPer7 probes targeted to indicated compartments to treatment of mtDAO-expressing cell lines with 4 mM D-Ala (black, wild type; red, PRDX1 KO; blue, PRDX2 KO; cell generated with the piggyBAC system). Cells were grown in glucose-containing medium. Solid line represents average, points colored in the lighter version of the respective color are the corresponding ratios measured in individual cells. **E.** Response of HyPer7 probes targeted to indicated compartments to treatment with 25 μM of exogenous H_2_O_2_. Indicated cell lines (black, wild type; red, PRDX1 KO; blue, PRDX2 KO) were grown in glucose-containing medium. Solid line represents average, points colored in the lighter version of the respective color are the corresponding ratios measured in individual cells. The numbers of cells per experiment as well as information on the statistical analysis for each dataset can be found in supplemental table **S6**. *, p ≤ 0.05, **, p ≤ 0.01, ***, p ≤ 0.001

Since HyPer7 steady state values were essentially unchanged between compartments and also the reactions to H_2_O_2_ were comparably small in the single knockout lines, we wondered whether PRDX1 and PRDX2 complemented for each other in the respective single knockouts. We thus generated a double knockout cell line lacking PRDX1 and PRDX2 (**Figures 8A, B**). Interestingly, also in this cell line many other antioxidative redox enzymes remained unchanged. Conversely, the HyPer7 steady state ratio differed strongly from the wild type in all compartments assessed including the mitochondrial matrix (**Figure 8C**). When we exposed the cell to an external bolus of H_2_O_2_, we observed a strong deflection of HyPer7 in all compartments, that appears to be limited by the dynamic range of the HyPer7 sensor (**Figure 8D**). HyPer7 in double knockout cells containing mtDAO was already at a highly oxidized steady state (without D-alanine addition) especially in the matrix and cytosol. Addition of D-alanine then led only to a minimal deflection because also here the sensor appeared to be limited by its dynamic range (**Figure 8E**).

**Figure 8.**
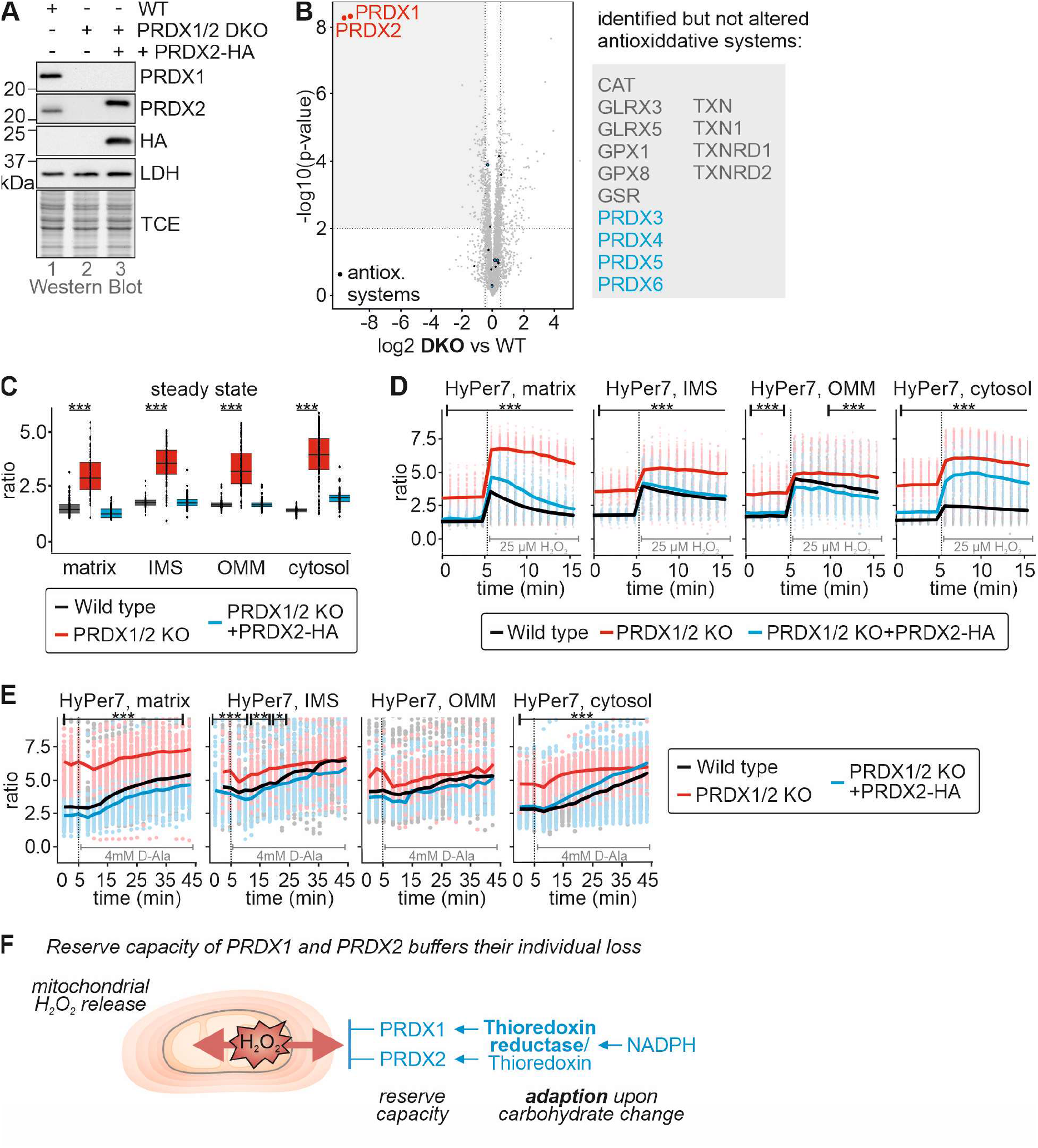
Peroxiredoxin 1 and 2 double knockout cells are strongly impaired in handling of mitochondrial H_2_O_2_ release. **A,B.** Peroxiredoxin levels in peroxiredoxin 1 and 2 double knockout cells (PRDX1/2 DKO). Lysates of the indicated cell lines grown in glucose-containing medium were analysed by immunoblot (**A**) and quantitative proteomics (**B**). **C.** HyPer7 steady state ratios of the indicated cell lines. HyPer7 probes were targeted to the indicated compartments of the respective cell lines. Cells were grown in glucose-containing medium. **D.** Response of HyPer7 probes targeted to indicated compartments to treatment with 25 μM of exogenous H_2_O_2_. Indicated cell lines (black, wild type; red, PRDX1/2 DKO) were grown in glucose-containing medium. Solid line represents average, points colored in the lighter version of the respective color are the corresponding ratios measured in individual cells. **E.** Response of HyPer7 probes targeted to indicated compartments to treatment of mtDAO-expressing cell lines with 4 mM D-Ala (black, wild type; red, PRDX1/2 DKO; cell generated with the piggyBAC system). Cells were grown in glucose-containing medium. Solid line represents average, points colored in the lighter version of the respective color are the corresponding ratios measured in individual cells. **F.** Model. *See discussion for details* The numbers of cells per experiment as well as information on the statistical analysis for each dataset can be found in supplemental table **S6**. Statistical analysis was only performed to compare the PRDX1/2 DKO with the wildtype. *, p ≤ 0.05, **, p ≤ 0.01, ***, p ≤ 0.001

Collectively, we demonstrate that cytosolic dithiol peroxiredoxins contribute to cytosolic H_2_O_2_ handling in HEK293 cells. They are present in amounts with sufficient reserve capacity so that either PRDX1 and PRDX2 can comfortably complement for the loss of the other enzyme, especially under unperturbed conditions, and that under conditions of increased H_2_O_2_ generation only the PRDX-regeneration thioredoxin system needs to become upregulated. The loss of both PRDX1 and 2 results in strong oxidation of cytosolic HyPer7 even under steady state conditions (**Figure 8F**).

## DISCUSSION

Here, using the ultra-sensitive, genetically encoded H_2_O_2_ sensor, HyPer7, targeted to different subcellular locations, we investigated the intracellular diffusion of mitochondrial H_2_O_2_ in single mammalian tissue culture cells. Our experiments yielded several interesting findings, including i) Peroxiredoxins strongly limit the intracellular diffusion of H_2_O_2_. ii) Peroxiredoxin activity is limited by the availability of thioredoxin reductase, the level of which varies between cell types, explaining previous contradictory observations. iii) Both H_2_O_2_ production and H_2_O_2_ removal capacity are modulated by changes in carbon metabolism, i.e. glucose vs. galactose. iv) Mitochondrial H_2_O_2_ does not reach the nucleus in detectable quantities, thereby raising important questions regarding the proposed models of mitochondrial H_2_O_2_ signaling. v) We observed a high degree of cell-to-cell heterogeneity in the ability to scavenge H_2_O_2_.

### Cytosolic peroxiredoxins limit amounts of mitochondrial H_2_O_2_ in the cytosol

We observed release of mitochondrial H_2_O_2_ using either antimycin A or DAO to induce mitochondrial H_2_O_2_ production. Using HyPer7 probes, localized to several different subcellular domains, we were able to show that mitochondrial H_2_O_2_ was largely confined to the proximity of mitochondria, as no response of PM or nuclear-localized probes was observed. In HEK293 cells we observed a small oxidation of the cytosolic HyPer7, which we did not observe in HeLa cells, consistent with previous observations. It is important to reiterate here, that we do not interpret our data as reflecting differences in the release of mitochondrial H_2_O_2_ between different cell types, particularly in the mtDAO expression studies. Instead, we believe that the difference reflects different cytosolic H_2_O_2_ scavenging capacities. The cytosolic HyPer7 probe is freely diffusible in the cytosol and will report on changes in the average H_2_O_2_ content of the whole cytosol. In HeLa cells, with very efficient H_2_O_2_ scavenging in the cytosol the diffusion of mitochondrial H_2_O_2_ will be strongly limited, leading to a steep H_2_O_2_ concentration gradient around the mitochondria. Likely, this region of cytosol is simply too small to affect the average cytosolic HyPer7 oxidation. On the contrary, in HEK293 cells, with less efficient cytosolic H_2_O_2_ scavenging, mitochondrial H_2_O_2_ will diffuse further into the cytosol, the H_2_O_2_ concentration gradient surrounding mitochondria will be less steep, the volume of cytosol with increased H_2_O_2_ levels will be higher and the average HyPer7 probe oxidation will give a detectable increase.

We demonstrate that diffusion of mitochondrial H_2_O_2_ in the cytosol is limited by PRDX1 and PRDX2. Both proteins are present in amounts that are high enough to handle much more H_2_O_2_ than is normally present in the cell. Whilst deletion of either one of these PRDXs did not induce any compensatory upregulation of the other PRDX in glucose-grown cells, we nonetheless observed that HyPer7 steady state oxidation remained comparable to the wildtype situation. Only upon rapid excess H_2_O_2_ generation/exposure did HyPer7 in the PRDX knockout cells become more oxidised than in the wildtype situation. Furthermore, when we deleted both PRDX together. the HyPer7 steady state became more oxidized; to an extent that hampered further dynamic analysis of mitochondrial H_2_O_2_ release. In summary, there appears to be considerable reserve capacity for H_2_O_2_ scavenging in terms of peroxiredoxin availability.

### Thioredoxin reductase availability is an important regulator of cytosolic H_2_O_2_ scavenging capacity

Despite the apparent high reserve capacity in terms of peroxiredoxin availability for H_2_O_2_ scavenging, we still observed considerable differences between different cell types in terms of H_2_O_2_ removal, determined by the ability of H_2_O_2_ to diffuse through the cytosol. Intriguingly, the level of thioredoxin reductase appears to be an important limiting factor for H_2_O_2_ scavenging. Comparing both HEK293 and HeLa cells and between glucose and galactose grown cells, we observed differences in thioredoxin reductase levels that correlated well with measured differences in H_2_O_2_ scavenging. Cells adapted to galactose exhibited increased protein amounts of cytosolic thioredoxin reductase (but not PRDXs). These cells also demonstrated an improved capacity to handle H_2_O_2_ thereby efficiently limiting the amounts of cytosolic H_2_O_2_ upon mitochondrial H_2_O_2_ generation. Collectively, our data obtained with genetically encoded fluorescent sensors are thereby in line with previous *in vitro* and in cell studies that reported that the thioredoxin system strongly impacts on PRDX activity and H_2_O_2_ dynamics in cells (Portillo-Ledesma, Randall et al., 2018, Toppo, Flohe et al., 2009). Our data also explain seemingly contradictory results in which in HeLa cells no impact of mitochondrial H_2_O_2_ release could be detected (Pak et al., 2020). In these cells, levels of thioredoxin reductase are higher compared to HEK293 cells essentially abolishing oxidation of cytosolic HyPer7 upon release of mitochondrial H_2_O_2_. Such differences might also explain differences in mitochondrial H_2_O_2_ signaling between tissues and during differentiation.

It is currently unclear why this should be so, although speculatively it could relate to conservation of NADPH. It is also unclear why peroxiredoxins are apparently present at much higher amounts than is necessary for H_2_O_2_ scavenging, although perhaps this relates to their other functions, for example as molecular chaperones (Rhee & Woo, 2020, Troussicot, Burmann et al., 2021). In summary, the rate-limiting step for cytosolic H_2_O_2_ handling and thus cytosolic detection of mitochondria-generated H_2_O_2_ appears to be in the reductive regeneration of PRDXs.

### Implications for mitochondrial H_2_O_2_ signaling and its crosstalk with metabolic adaptations

Adaptation of cells to galactose rebalances cellular H_2_O_2_ dynamics. On the one hand, it improved H_2_O_2_ handling by strengthening the reductive regeneration of PRDXs; on the other hand, it also increased the mitochondrial capacity to generate H_2_O_2_. This led to an unchanged HyPer7 steady state between both carbon sources. However, metabolic adaptations might also offer a temporal window of opportunity for redox signaling as initial flux through the respiratory chain might result in increased generation of H_2_O_2_ that is initially not matched by upregulation of antioxidative systems.

Due to the steep gradients of H_2_O_2_ around mitochondria, signaling by mitochondrial H_2_O_2_ likely takes place in close proximity to mitochondria. Such a confined nature of mitochondrial H_2_O_2_ signaling could enable signaling pathways originating from mitochondrial subpopulations or specific parts of the mitochondrial network within a cell, e.g. reflecting intracellular heterogeneities in the mitochondrial network with respect to the membrane potential of respiratory chain activity. Our data also support the use of PRDX-mediated pathways for mitochondrial H_2_O_2_ signaling (Sobotta, Liou et al., 2015, Stocker, Maurer et al., 2018). The detected cytosolic levels of H_2_O_2_ upon mitochondrial H_2_O_2_ generation are high enough to oxidize HyPer7, and our PRDX knockout data indicate that endogenous PRDXs can efficiently compete with HyPer7 for H_2_O_2_, implying that endogenous PRDXs can efficiently sense the released H_2_O_2_.

In our single cell measurements, we observed substantial intercellular differences in HyPer7 responses. Such heterogeneities are often masked when assessing population averages. We currently can only speculate on their origin, e.g. that they represent responses of cells at different stages of the cell cycle or cells that maintain different fluxes through pathways providing reducing equivalents for H_2_O_2_ detoxification, or cells in which mitochondria exhibit different respiratory chain activities. Using single cell multiplexing approaches to e.g. concomitantly assess H_2_O_2_ dynamics and cell cycle progression or the mitochondrial membrane potential might in future studies help to mechanistically explain the observed heterogeneities.

## MATERIALS AND METHODS

### Plasmids and Cell lines

For plasmids, cell lines, primers, CRISPR guides and antibodies used in this study, see **Tables S1-S5**. All cell lines were cultivated using Dulbecco’s modified Eagle’s medium (DMEM) complete containing 4.5 g/l glucose, 10 % fetal bovine serum (FCS), and 500μg/ml penicillin/streptomycin at 37 °C under 5% CO_2_.

### Generation of Peroxiredoxin knockout cells

Guide RNA sequences targeting human PRDX1 or PRDX2 were cloned into the pSpCas9(BB)-2A-GFP (PX458) vector, which was a gift from Feng Zhang (Addgene plasmid # 48138) (Ran et al.,2013). HEK Flp-In™ T-REx™-293 cells were transfected using PEI. After 24h, FACS sorting was used to collect GFP-positive cells. Single cells were seeded into 96-well plates. Clones were screened using western blot. For complementation, the Flp-In™T-REx™ system (Invitrogen) was used. For selection of positive clones, DMEM complete containing 100 μg/ml hygromycin and 10 μg/ml blasticidin was used. Prior to experiments, the expression of stable cell lines was induced for at least 16 h with 1 μg/ml doxycycline. To express matrix targeted DAO in the PRDX knock-out cell lines, the Su9-DAO construct was cloned into the cumate-inducible PB-CuO-MCS-IRES-GFP-EF1-CymR-Purovector (System Biosciences; PiggyBac system). Positive clones were selected with DMEM complete containing 2μg/ml puromycin and expression of the cell line was induced with 30 μg/ml cumate for at least 16h before the experiment.

### Steady state protein levels

HEK239 MOCK cells grown in glucose or galactose DMEM complete were harvested in 1x Laemmli buffer (2% SDS, 60 mM Tris-HCl pH 6.8, 10% glycerol,0.0025% bromophenol blue), boiled at 96°C for 5 min and subsequently analyzed by SDS-PAGE and western blot. As loading control, 2,2-trichloroethanol (TCE) was added to the SDS gel to visualize protein levels.

### Immunofluorescence

HeLa cells were seeded onto poly-L-lysine coated cover slips in DMEM complete medium. After 24 h, cells were transfected with the respective HyPer7 sensor using PEI (polyethylenimine). After 48 h, cells were fixed using 4 % paraformaldehyde for 15 min, blocked with BSA-blocking buffer (10 mM HEPES, 3 % BSA, 0.3 % triton-X-100) for 1h and incubated with self-made roGFP2 primary antibody or FLAG antibody for 1 h at RT. Subsequently, incubation with secondary antibody STAR 635P goat anti-rabbit (abberior) or Alexa Fluor 488 goat anti-muse (invitrogen) for 1 h at RT was followed by conserving the samples with mounting medium (30% glycerol, 12% polyvinyl alcohol, 60 mM TRIS, 2.5% 1,4-diazabicyclo-2,2,2-octan).

STED microscopy for the HyPer7 localization was performed on a Leica TCS SP8 gSTED 3X system (Leica Microsystems) using a 93x glycerin objective with a numerical aperture of 1.3 (HC PL APO CS2 93x/1.30 GLYC, Leica Microsystems). For gated STED a pulsed white light laser at 633 nm and a 775 nm STED depletion laser was used. Images were deconvolved using the software Huygens Essential (Scientific Volume Imaging).

### Quantitative label-free proteomics

Cells were seeded in 6-well plates (n=4). After reaching confluence, cells were washed once with PBS, scratched off in 1ml PBS and transferred in a 1.5 mml reaction tube. After 5 min of centrifugation at 500 g, supernatant was removed, and lysis buffer was added (4 % SDS in PBS containing protease inhibitor). Samples were sonicated 20 times (60×60) and boiled at 96 °C for 5 min to precipitate proteins, fourfold volume of ice-cold acetone was added, and samples were frozen at −80 °C. After thawing, samples were centrifuged for 15 min at 16,000 g, supernatant was removed, and samples were washed twice with 500 μl ice-cold acetone. The pellet was air dried. In-solution digest and stage tipping were performed according to the protocols of the proteomics facility from CECAD (https://www.proteomics-cologne.com/protocols). All samples were analyzed on a Q Exactive Plus Orbitrap (Thermo Scientific) mass spectrometer that was coupled to an EASY nLC‘‘LC’’ (Thermo Scientific). Peptides were loaded with solvent A (0.1% formic acid in water) onto an in-house packed analytical column (‘containing 2 layers of SDB-RPS disc). Peptides were chromatographically separated at a constant flow rate of 250 nL/min using the following gradient: 3%–5% solvent B (0.1% formic acid in 80% acetonitrile) within 1.0 min, 5%–30% solvent B within 91.0 min, 30%–50% solvent B within 17.0 min, 50%–95% solvent B within 1.0 min, followed by washing and column equilibration. The mass spectrometer was operated in data-independent acquisition mode. The MS1 scan was acquired from400-1220 m/z at a resolution of 140,000. MSMS scans were acquired for 10 DIA windows at a resolution of 35,000. The AGC target was set to 3e6 charges. The default charge state for the MS2 was set to 4. Stepped normalized collision energy was set to 23.5%,26%, 28.5%. The MSMS spectra were acquired in profile mode.

### HyPer7 and roGFP2-Prx1 measurements

4,000 cells/well were seeded in 100 μl complete DMEM on a poly-L-lysine coated 96-well plate (μclear, GreinerBio). After 24 h, cells were transfected with plasmids containing sensors using PEI. After 48 h, DMEM was replace by minimal medium (140 mm NaCl, 5 mm KCl, 1 mm MgCl_2_, 2 mm CaCl_2_, 20 mm Hepes, 10 mm glucose, pH set to 7.4 with NaOH) and the 96-well plate was incubated inside Cytation3 (BioTek) for 40 min at 37°C and 5 % CO_2_. For all measurements a 10x in-air microscope and the BioTek LED filter cubes 390+/-15 nm and 467+/-15 nm were used. For all measurements, 5 min of steady state were measured. Two kinds of H_2_O_2_ titration were performed. In one, different volumes (20, 40 and 80 μl) of a solution with a very low H_2_O_2_ concentration were added, each after 20 min, to the same well sequentially, resulting in concentration of ~ 2, 4 and 8 μM H_2_O_2_ in the well. Finally, DTT was added as control (50 μl, resulting in 20 mM). In the other, 12.5 μM H_2_O_2_ were added to single wells and measured for 20 min. DTT was added as control as well. To investigate H_2_O_2_ in cell grown in different carbon source, they were grown for at least 1 day in the respective carbon source before conducting the experiment. To investigate to production of H_2_O_2_ upon inhibition of the respiratory chain, 30 μl of MM containing Antimycine A (20 μM f.c.) were added to different wells and measured for 50 min.

For a D-Alanine titration, after 5 min steady state, 30 μl of MM containing D- or L-Ala resulting in 1, 2, 4, or 8 mM D-Alanine or 8 mM L-Alanine were added to different wells and measured for 50-60 min. To investigate if inhibition of TXNRD1 influence H_2_O_2_ dynamics, to HEK293 matrix-DAO cells grown in galactose 2 mM D- or L-Ala were added combined with or without 1 μM auranofin after 5 min of steady state measurements. Sensor oxidation was followed for 50 min.

For the hypoxia/reoxygenation experiments, cells were prepared as for the other experiments. 10 min of steady state were measured at 20 % O_2_. Then hypoxia was started and reached 1 % O_2_ after 30 min. After 4 h at 1 % O_2_, reoxygenation was started and 20 % O_2_ were reached after 1h. During the whole time, the measurements were conducted every 15 min.

### Data analysis, quantification and statistical analysis

The acquired images for each experiment were analyzed using the program RRA (“redox ratio analysis”, (Fricker, 2016)). With this program, images were aligned, filtered, background-subtracted and the intensity for both channels as well as the resulting ratio (500/400 or 405/488) was calculated and saved as an excel file. Using R, these excel files were further analyzed, the mean was calculated, figures prepared, and statistics performed. For Cytation3 measurements, the data was first analyzed for normal distribution using a Shapiro-Wilk-test. As most of the data were not normally distributed, instead of a t-test, a Wilcoxon/Mann-Whitney-U-test was performed and samples were compared in pairs. Western blot signals were quantified using Image Lab 5.2.1 (*Bio-Rad*). Error bars represent standard deviation.

## ACKNOWLEDGEMENTS

The Deutsche Forschungsgemeinschaft (DFG) funds research in the Laboratory of JR (RI2150/2-2 – project number 251546152, RI2150/5-1 – project number 435235019, CRC1218 / TP B02 – project number 269925409, and RTG2550/1 – project number 411422114). VVB was supported by the grant 075-15-2019-1789 from the Ministry of Science and Higher Education of the Russian Federation. We thank the CECAD Proteome and Imaging Facilities for the provision of instrumentation, training, and technical support.

## AUTHOR CONTRIBUTIONS

JR and MNH designed the study and planned experiments. MNH, LJ, KL, LM, TM performed experiments. JR, LJ and MNH analysed data. MF wrote the RRA software and trained MNH in its use. VVB provided the HyPer7 plasmid before publication and provided critical input into the planning of the study. BM provided critical input into the planning of the study. JR, BM, VVB and MNH wrote the manuscript with critical input from all authors.

